# Spatial lung niches shape *Pseudomonas aeruginosa* persistence

**DOI:** 10.64898/2026.07.01.735213

**Authors:** Cecilia Kyany’a, Duy Pham, Nana-Jane Chipampe, Phumrapee Boonklang, Louise Ellison, Andrew J. Balmer, Catherine Tudor, Jasmine Halliwell, Hon Man Chan, Vanessa Brito Pereira, Benjamin Rumney, Holly Anderson, Shevaniee Umamaheswaran, Mia Franulovic, Terry K. Judah, Sam Dougan, Minal Patel, Omer Bayraktar, Hannah Wong, Kenny Roberts, Sarah A. Teichman, R. Andres Floto, Josephine M. Bryant

## Abstract

Chronic *Pseudomonas aeruginosa* lung infection is a major cause of morbidity and mortality in people with pre-existing lung disease. Once established, infection is rarely eradicated and often persists despite prolonged antimicrobial therapy, but the underlying mechanisms remain poorly understood. To define how *P. aeruginosa* occupies and adapts to diseased lung tissue, we applied host–pathogen spatial transcriptomics to profile over 23 million lung cells from explanted and resected lungs from people with Cystic Fibrosis (CF) and chronic obstructive pulmonary disease (COPD), mapping bacterial niches and transcriptional states *in situ*.

We found that *P. aeruginosa* adopted distinct niche-linked states across chronically infected human lung tissue. Bacterial burden was highest in airway lumens, but bacteria also occupied submucosal glands, parenchyma and, unexpectedly, blood vessel lumens in CF tissue. Within airway lumens, two coupled host–pathogen states emerged: an alginate-rich biofilm-like state linked to PI3-positive neutrophil inflammation and host chemical-sensing programmes; and a motile, quorum-sensing state with activated type 6 secretion linked to human ciliary stress, epithelial remodelling and proteolytic injury. Intravascular bacteria co-localised with neutrophils, fibrin and vascular-remodelling signatures, suggesting local breach of barrier integrity and subsequent immune containment. *P. aeruginosa* also displayed disease-specific cellular associations, with neutrophil-dominated interactions in CF and greater association with macrophages and dendritic cells in COPD.

**Together, these data reveal chronic *P. aeruginosa* infection as a spatially partitioned ecosystem in which anatomical microenvironments impose distinct bacterial lifestyles and host inflammatory states.** This niche-resolved framework helps explain how persistent infection can diversify within a single lung and suggests that eradication therapies may need to target multiple anatomical and cellular niches to be effective.

## Main

Microenvironmental and cellular diversity within the human body exerts powerful selective pressures on invading pathogens, shaping their evolution, persistence, and clinical impact. *P. aeruginosa* is a major opportunistic human pathogen, killing more than 800,000 people annually of which almost half are associated with antimicrobial resistance, and is particularly notable for its adaptability^1^, establishing infections in a wide range of human tissues^2–5^. In the lung, *P. aeruginosa* infections can persist for years despite prolonged antibiotic exposure and ongoing immune activation. The mechanisms underlying this persistence remain unresolved, but likely derive from this species’ capacity for phenotypic plasticity^6^ and mutational adaptation^1,7,8,9^, enabling survival in heterogeneous niches.

The lung itself comprises diverse ecological niches with differing cellular architecture, immune cell populations^10,11^, and biochemical properties. While *P. aeruginosa* has long been regarded as primarily an extracellular pathogen, persisting in sputum and biofilms of the bronchial airways^12^ emerging evidence suggests that it can also survive intracellularly within goblet cells^13^, epithelial cells^14,15,16^ , and macrophages^4^. However, *in situ* evidence for such intracellular persistence in human lung tissue remains very limited.

These observations therefore raise the possibility that chronic persistence is not explained solely by biofilm-mediated antibiotic tolerance in the airway lumen, but by microenvironment-driven switching between distinct bacterial physiological states. We hypothesised that *P. aeruginosa* may exploit multiple lung microenvironments to optimise survival. To investigate this, we applied dual host and pathogen spatial transcriptomics to explanted lung tissue from people with Cystic Fibrosis (CF) and COPD, providing a unique opportunity to study the spatial distribution and adaptive strategies of *P. aeruginosa* directly in human tissue.

### *P. aeruginosa* dominates polymicrobially infected lung tissue and expresses lung-adaptive programmes

We analysed explanted or resected lung tissue (formaldehyde-fixed and paraffin-embedded; FFPE) from 13 adults with chronic *P. aeruginosa* lung infection (11 with CF; 2 with COPD, Supplementary table 1). We first performed bulk RNAseq on 52 tissue sections from different lung lobes to detect both host and bacterial RNA. For the human transcripts, we employed *in silico* “cytometry” to infer cell composition for each sample (using immune (LM22) and extended (Human Cell Atlas) databases; Fig. 1a). We observed disease specific patterns of cellular composition, with a higher abundance of neutrophils (adjusted p-value = 0.0005) and B cells (adjusted p-value = 0.0037) in CF and higher proportions of interstitial macrophages (adjusted p-value = 0.00081) in COPD.

**Figure 1.**
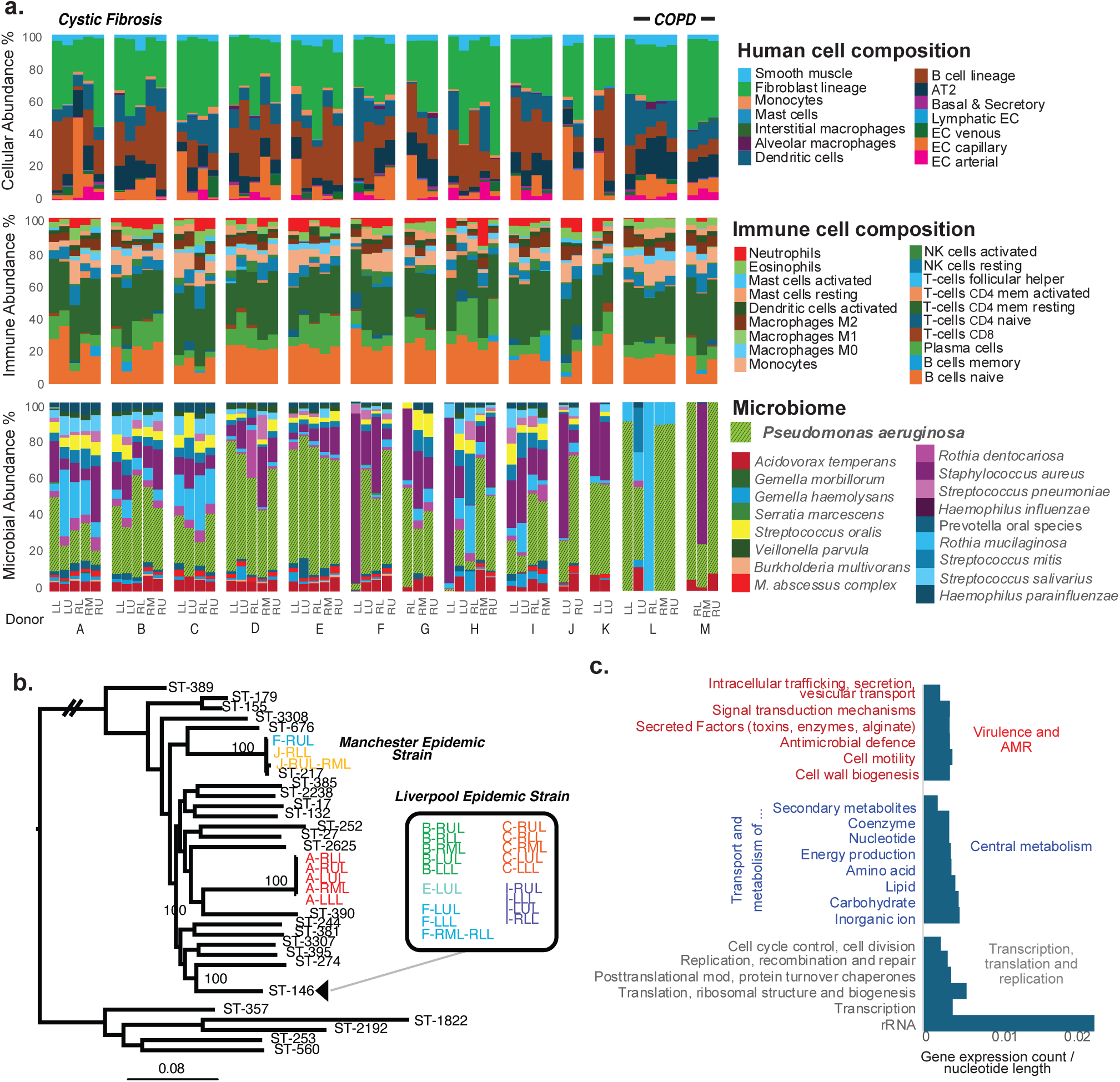
Bulk transcriptomic and phylogenetic analysis of *P. aeruginosa* from CF infected lung tissue. (a) We performed bulk RNAseq on 52 FFPE tissue sections and employed in silico “cytometry” to infer lung cell (Human Cell Atlas) and immune (LM22) cell composition. Bulk metatranscriptomics was performed and used to infer the microbiome diversity present in the tissue in donors A-M. Due to the possible presence of contaminant reads, the bar plot is restricted to known bacterial commensal and pathogen species previously associated with CF and COPD. *P. aeruginosa* transcripts were detected in all of the donors. LL=left lower lobe, LU=left upper lobe, RL=right lower lobe, RM = right middle lobe, RU=right upper lobe. (b) Maximum likelihood phylogeny of *P. aeruginosa* genomes. A genome-wide *P. aeruginosa* bait panel was used to enrich for *P. aeruginosa* DNA extracted from the FFPE lung tissue. Phylogenetically informative SNPs could be identified with sufficient coverage in 28 tissue blocks from 7 of the donors. In two cases the reads from multiple blocks were merged (J and F) to enable accurate phylogenetic placement. Genomes from STs common in CF were included for context, and the support for key nodes from 100 bootstrap replicates are indicated. One branch was shortened for illustration purposes (indicated by double lines). 6/7 patients were found to be infected with either the Liverpool or Manchester Epidemic Strains. (c) Transcripts specific to *P. aeruginosa* were extracted from the metatranscriptomic data and mapped to gene functions. Sparse read coverage did not permit normal genome-wide differential expression analysis, but instead we counted genes where atleast one read-pair mapped and divided by the the gene length as there is greater opportunity to detect a transcript for longer genes. The genes were assigned to COG categories. As expected the most highly expressed transcripts were ribosomal RNAs followed by protein coding genes associated with transcriptional, translational, replication and core metabolism functions. We also detected transcripts predicted to be associated with the secretion of toxins and signalling which may be involved in interactions with human lung cells and niche adaptation.

For bacterial transcripts, we assessed the level of microbial diversity across samples. After removal of contaminant reads, we found that *P. aeruginosa* was the dominant bacterium in 73% (38/52) of sections, with remaining reads assigned to other pathogens and commensals commonly found in chronic lung disease (Fig. 1a). Targeted DNA bait capture enabled recovery of *P. aeruginosa* genomes from tissue and showed that, in all but one donor, the same strain was present across all sampled anatomical regions within an individual (Fig. 1b). Six of the seven CF donors with sufficient read depth, were found to be infected with epidemic CF-associated clones known to infect a large proportion of adults with CF in the UK^17^. Analysis of *P. aeruginosa* transcripts revealed high expression of housekeeping genes and genes encoding motility machinery and secreted virulence factors that have previously been shown to be important for *in vivo* persistence (Fig. 1c).

Guided by these findings, we designed a *P. aeruginosa*-specific probe panel for spatial profiling on 10x Genomics Xenium and Visium platforms, targeting genes involved in core cellular function (16S rRNA, 23S rRNA, *groEL*), motility (*rpoN*), alginate biosynthesis (*algE*), toxin secretion (*exoT, exoS, hcp1, toxA,* and *pcrV*), quorum sensing (*lasI* and *lasR*) and a predicted contact-dependent inhibition gene (PA2462) which was particularly highly expressed in our data (detected in 23/52 sections). Probe sequences are listed in Supplementary Table 3-4.

### Spatial profiling identifies large airways as the principal *P. aeruginosa* niche

For spatial profiling, we prioritised patients with high *P. aeruginosa* abundance based on bulk RNA-seq. Twenty-eight tissue sections (four per subject) from five CF and two COPD donors were selected for profiling on the 10x Genomics Xenium In Situ platform, including a sample from every available lobe from each donor. Using a custom *P. aeruginosa* probe panel targeting the aforementioned 13 bacterial genes alongside the Xenium Human Lung Panel v1 (289 genes), we detected both *P. aeruginosa* rRNA and mRNA transcripts throughout tissue sections (Fig. 2a, Extended data Fig. 1a,b). In total, we profiled more than 23 million cells and 15.4 million *P. aeruginosa* transcripts across chronically infected human lung tissue, generating a large-scale spatial resource for studying host–pathogen interactions in advanced lung disease.

**Figure 2:**
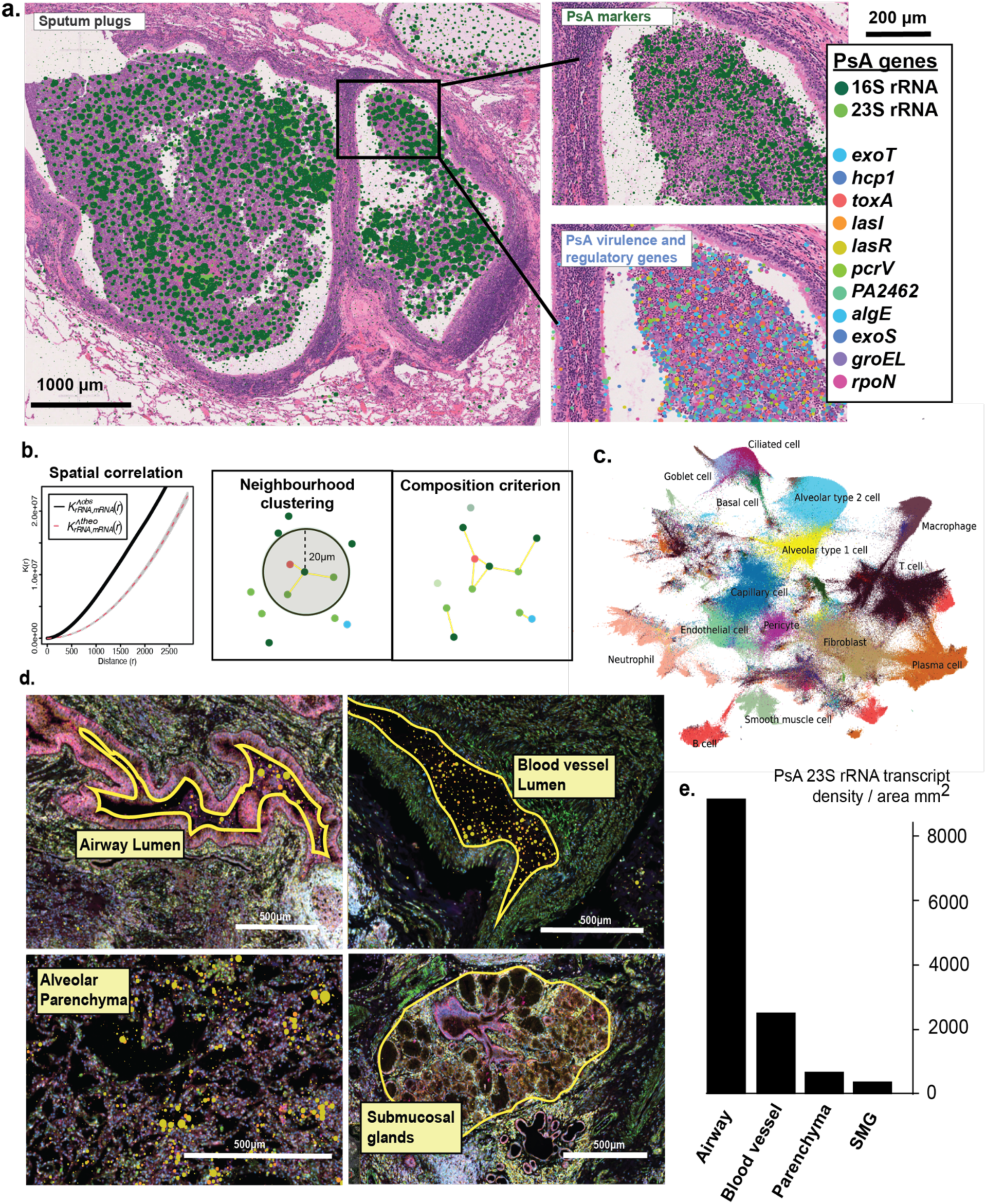
Detection of *P. aeruginosa* transcripts on Xenium In Situ Spatial Imaging. PsA = *P. aeruginosa* (a) Representative Xenium overlay with H&E showing detection of *P. aeruginosa* transcripts in the airway lumen from donor B RML. The highlighted inset shows detection of *P. aeruginosa* rRNA (top) and mRNA (bottom) transcripts. (b) Ripley K spatial correlation to filter out transcripts without spatial correlation and Infographic showing Ripley K spatial correlation of *P. aeruginosa*. We considered only mRNA and rRNA transcripts with spatial correlation within a 20µm radius for downstream analysis. (c) UMAP for 1 million subsampled cells using rapid-sc (d) Representative Xenium image (Xenium Explorer vs4) of anatomical regions of interest. Yellow line show boundaries and *P. aeruginosa* transcripts are indicated by coloured spots. (e) Bar plot of *P. aeruginosa* 23S RNA across anatomical regions of interest. The highest *P. aeruginosa* densities were detected within the airway lumens. SMG= submucosal glands.

To distinguish true bacterial signal from background noise, we applied Ripley’s K spatial correlation and retained only co-localised *P. aeruginosa* transcripts arising from distinct gene targets and containing at least one rRNA transcript (Fig. 2b). Since *P. aeruginosa* has been shown to exist in micro-colonies^18^ and biofilms^19^ during chronic lung infections, we considered spatially correlated transcripts within a 20µm radius (Fig. 2b). This filtering retained a median of 86% of transcripts per section (range 42–99%, Extended data Fig. 1c), corresponding to 15.2 million spatially correlated transcripts overall of which the majority (96.3%) were rRNA. *P. aeruginosa* abundance measured by bulk RNAseq correlated with abundance detected by Xenium profiling (Spearman’s ρ = 0.418, p = 0.028; Extended data Fig1d). Human cells were segmented and assigned to cell types using established workflows, identifying the expected features of inflamed chronic lung tissue, including abundant immune-cell infiltration (Fig. 2c).

*P. aeruginosa* transcripts were unevenly distributed across tissue sections, with very dense signals in airway lumens, frequently corresponding to sputum plugs (Extended data Fig. 1a and Fig. 2a). However, in CF patients, additional signal was detected within submucosal glands (SMG), parenchyma, and blood vessel lumens. To quantify bacterial distribution across anatomical compartments, we manually annotated 312 regions across the 17 slides from the 5 CF donors (Fig. 2d). Quantification across annotated regions confirmed that *P. aeruginosa* density was highest in airway lumens. Unexpectedly, blood vessel lumens showed the second highest bacterial density (Fig. 2e), with 62 of 115 annotated regions containing two or more spatially correlated 23S rRNA transcripts (Supplementary table 5), indicating a previously unrecognized vascular niche in CF lung tissue.

### Intravascular localisation of *P. aeruginosa* in CF lung tissue

To further investigate the presence of *P. aeruginosa* within blood vessels, we performed matched spatial profiling of the same tissue sections using Xenium imaging-based and Visium sequencing-based technologies with custom *P. aeruginosa* probe panels. We detected concordant signal between the two platforms in all cases suggesting this was not an artefact of the technology (Fig. 3a, Extended data Fig. 1a). We validated our transcription-based finding using immunofluorescence with a polyclonal antibody against *P. aeruginosa* and observed dense signal within blood vessel lumens (Fig. 3b, Extended Fig 2a), supporting our initial observation. Intravascular signal was observed across multiple tissue sections across multiple donors (Fig. 3c).

**Figure 3:**
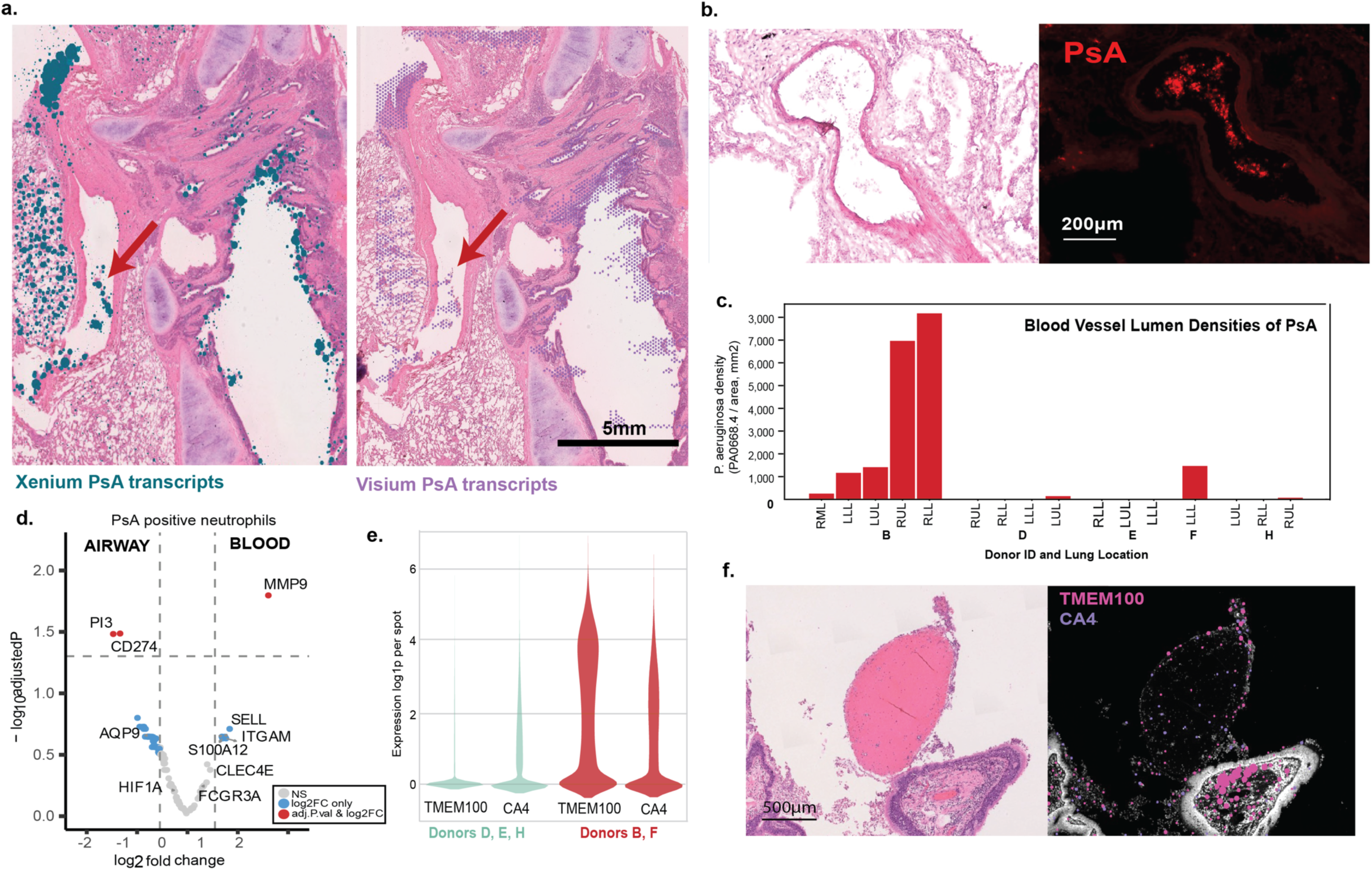
Detection of *P. aeruginosa* transcripts within blood vessel lumens. PsA = *P. aeruginosa* (a) Representative tissue section image from Donor B RML profiled on both Xenium and Visium showing location of *P. aeruginosa* transcripts within the blood vessel lumen (red arrowhead). (b) Immunofluorescence image showing *P. aeruginosa* within blood vessel of Donor D. Left image is H&E showing blood vessel. Right image is IF showing *P. aeruginosa (*in red) within the blood vessel lumen. (c) Bar plot of *P. aeruginosa* rRNA 23S transcript density across blood vessel lumens annotated across the five CF donors. (d) Differential gene expression analysis between *P. aeruginosa* positive airway neutrophils and *P. aeruginosa* positive blood neutrophils. Genes with significant adjusted p-values (<=0.05) and log2 fold change >=0.58 (∼1.5-fold increase) are shown in red. (e) Expression of vascularisation associated genes, *TMEM100* and *CA4*, per Visium spot for donors with significant blood colonisation (red) and those without (green). Expression was significantly higher in the patients with blood colonisation (mixed linear model *TMEM100*: p = 8.47e-6; *CA4*: p=2.04e-2). (f) Colocalisation of *TMEM100* and *CA4* expression in the airway subepithelial region (mucosa and submucosa) adjacent to a blood and fibrin aggregate in the airway of donor B RLL. Data generated on Xenium Prime 5k and visualised as H&E stain (left) and Xenium morphology image (right). Image made using Xenium Explorer vs4.

We considered the possibility that intravascular bacterial signal could reflect tissue handling artefacts during pathological processing. If so, we would expect immune cells within blood vessel lumens to exhibit gene expression profiles similar to airway-associated cells. We found that, in both compartments, *P. aeruginosa* signal spatially co-localised with neutrophils. However, comparison of infected neutrophils from blood vessel and airway lumens revealed distinct transcriptional programmes. Intravascular *P. aeruginosa*-positive neutrophils had significantly higher expression of *MMP9*, whereas airway-associated neutrophils expressed higher levels of *PI3* (elafin) (Fig. 3d) and these differences were reproducible in bacteria-negative cells (Extended data Fig. 2b). These differences indicate that *P. aeruginosa*-associated neutrophils within blood vessels are unlikely to have originated from airway contamination and represent a distinct cellular population.

All five CF donors had detectable *P. aeruginosa* within blood vessel lumens, although vascular bacterial density varied markedly between individuals (Fig. 3c; Kruskal–Wallis H = 12.83, p = 0.012). Donor B, who had the highest vascular bacterial burden across all five profiled lung regions, had a clinical history of persistent haemoptysis, supporting a link between intravascular *P. aeruginosa* localisation and airway bleeding. Hypervascularisation and fibrin deposition were common histopathological features across CF tissues (Extended data Fig. 2c) and in several cases *P. aeruginosa* signal was embedded within fibrin deposits (Extended data Fig. 2d). Analysis of the Visium data which provided broader transcriptomic coverage at lower spatial resolution, identified significant spatial co-localisation between *P. aeruginosa* and expression of the vascular-associated genes *TMEM100* and *CA4* (Extended data Fig. 2e). These genes previously implicated in neovascularisation had significantly higher levels of expression in patients with intravascular *P. aeruginosa* (Fig. 3e,f). These findings implicate vascular remodelling and fibrin deposition in the intravascular localisation of *P. aeruginosa*.

### Airway *P. aeruginosa* states are coupled to distinct host microenvironments

To determine whether bacterial transcription varied by anatomical context, we compared *P. aeruginosa* gene expression across the annotated regions from CF donors. Principal component analysis showed that bacterial expression profiles from alveolar parenchyma were highly variable across samples, likely reflecting the structural heterogeneity of these regions. By contrast, expression profiles from submucosal glands, blood vessel lumens, and airway compartments clustered more tightly by anatomical site, indicating niche-specific bacterial transcriptional states (Fig. 4a). Differential expression analysis, adjusted for donor ID, identified significantly higher expression of *toxA* in submucosal glands compared with all other compartments (adjusted p-value: 1.54E-08, log2 fold change: 1.02. Extended data Fig. 3a).

**Figure 4:**
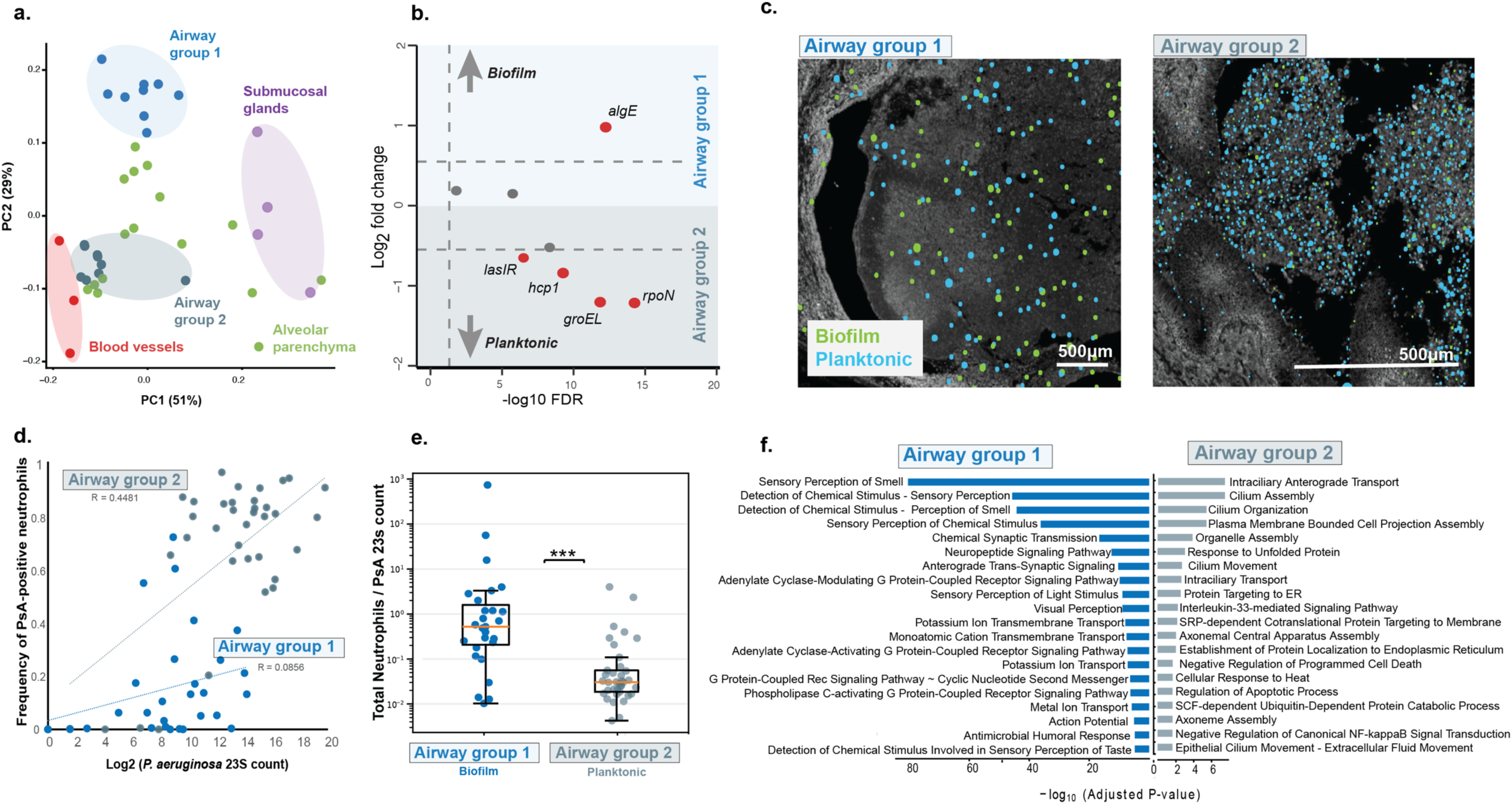
Anatomical niche defines distinct *P. aeruginosa* transcriptional states in CF lung tissue. PsA = *P. aeruginosa.* Annotated anatomical regions were manually defined and *P. aeruginosa* transcript counts extracted. Analyses included only CF donors, as insufficient anatomical regions were identifiable across COPD tissue sections. a) *P. aeruginosa* transcript counts from each annotated region were aggregated by tissue section, normalised to total bacterial mRNA abundance, and visualised by principal component analysis. Regions with fewer than 60 total transcripts were excluded. b) Volcano plot showing differential *P. aeruginosa* gene expression between airway group 1 and airway group 2. Red dots indicate genes that had greater than 1.5x fold change and were statistically significant. c) Example Xenium images made using Xenium Explorer vs4 from airways identified in donor E LUL (l) and donor B LLL (r) where green dots indicate biofilm related transcripts (*algE*) and blue dots indicate planktonic related transcripts (*hcp1, groEL, rpoN, lasI, lasR*). d) Neutrophils from annotated airway lumens, were classified as *P. aeruginosa* -positive (≥2 bacterial transcripts) or negative. Each point represents an individual airway lumen. Linear regression analysis identified a significant positive correlation between bacterial burden and neutrophil positivity in airway group 2 (p = 0.019), but not airway group 1. e) Box-and-whisker plot showing neutrophil abundance across airway lumens from airway groups 1 and 2. Statistical significance was determined using a two-sided Mann–Whitney U-test. f) Top 20 GO terms statistically associated with either airway group 1 or airway group 2 identified using differential expression analysis (p < 0.05 and log2FC > 3) and subsequent enrichment analysis, between airway (sputum) niches defined by applying CellCharter to Visium data.

Within airway lumens, two distinct *P. aeruginosa* transcriptional states were observed (Fig. 4a,b): Group 1 comprised all airways examined from donors E and H; Group 2 comprised airways from donors B and F; while both airway groups were seen in Donor D (Extended data Fig. 3b). Although broadly split by donor, this pattern of clustering was not explained by underlying *P. aeruginosa* genetic diversity, as donors B, E and F were all infected with the same epidemic strain (Fig 1b). Similarly, this separation was not accounted for by recorded comorbidities or demographic factors (Supplementary Table 1).

Differential expression analysis between airway groups identified significantly higher *algE* expression in group 1, consistent with alginate production and a biofilm-associated state^20^. In contrast, group 2 showed increased expression of *lasI/lasR, rpoN, groEL,* and *hcp1*, corresponding to quorum sensing, motility, growth, and type VI secretion system activity, respectively (Fig. 4b).

This profile is consistent with a planktonic, active-growth lifestyle^21^. Notably, airway group 2 transcriptional profiles closely resembled those observed in blood vessel lumens (Fig. 4a).

Previous studies have established that biofilm-associated *P. aeruginosa* resist phagocytosis^22^ trigger neutrophil swarming^23^ and can immobilise recruited neutrophils within the biofilm matrix ^24^ . We therefore tested whether the two airway transcriptional states differed in their neutrophil interactions. Group 2 airways showed a significant positive relationship between *P. aeruginosa* abundance and proportion of neutrophils positive for *P. aeruginosa* signal (P < 0.001), whereas Group 1 did not (Fig. 4d). In contrast, Group 1 had a substantially higher neutrophil burden per bacterial transcript than group 2 (median 0.52, range 0.21–1.59, versus median 0.03, range 0.019–0.056 neutrophils per *P. aeruginosa* 23S rRNA transcript; two-sided Mann–Whitney P < 0.0001; Fig. 4e). These data suggest two distinct airway states: a neutrophil-rich, biofilm-associated state in group 1, where bacteria appear embedded within dense inflammatory foci, and a more dispersed planktonic-like state in group 2, where neutrophil association scales with bacterial burden.

We next asked whether these bacterial airway states were associated with distinct host microenvironmental responses. Airway group 1, characterised by a biofilm-associated bacterial profile, showed strong enrichment of host genes involved in sensory perception of chemicals (Fig. 4f). This included *TAS2R38* (adjusted p value = 1.79e-73), a bitter taste receptor implicated in airway detection of bacterial products and host responses to *P. aeruginosa*^25,26^. These data suggest that the non-motile, biofilm-associated state is linked to a host programme of chemical sensing, potentially reflecting local detection of bacterial metabolites within chronically infected sputum.

For airway group 2 however, characterised by a faster-growing, motile and planktonic-like bacterial profile, was enriched for host programmes related to ciliary movement, epithelial repair and tissue remodelling, together with IL-33-associated inflammation. We also observed enrichment of unfolded-protein response and protein-catabolism pathways, consistent with epithelial stress and mucosal barrier damage. Such damage could release host-derived proteins and nutrients into the airway lumen, creating a feed-forward niche that supports bacterial growth and inflammatory activation.

Together, these findings suggest that airway *P. aeruginosa* exists in at least two coupled host–pathogen states: a biofilm-associated state linked to chemical sensing and chronic containment, and an active-growth state linked to epithelial injury, ciliary stress and inflammatory remodelling. This state structure may provide a spatial mechanism for transitions between stable chronic infection and exacerbation-like tissue damage.

### Distinct immune cell populations harbour transcriptionally distinct *P. aeruginosa*

To determine whether *P. aeruginosa* exhibited heterogeneous cellular associations, we performed cell assignment across all Xenium datasets and identified cell types significantly enriched for bacterial transcripts relative to other cells from the same donors (Fig. 5a). Analyses were restricted to cell types represented by at least 1,000 cells across all donors, with CellAssign confidence scores ≥0.9. Cells containing ≥2 spatially correlated *P. aeruginosa* transcripts were classified as bacterial-positive.

**Figure 5:**
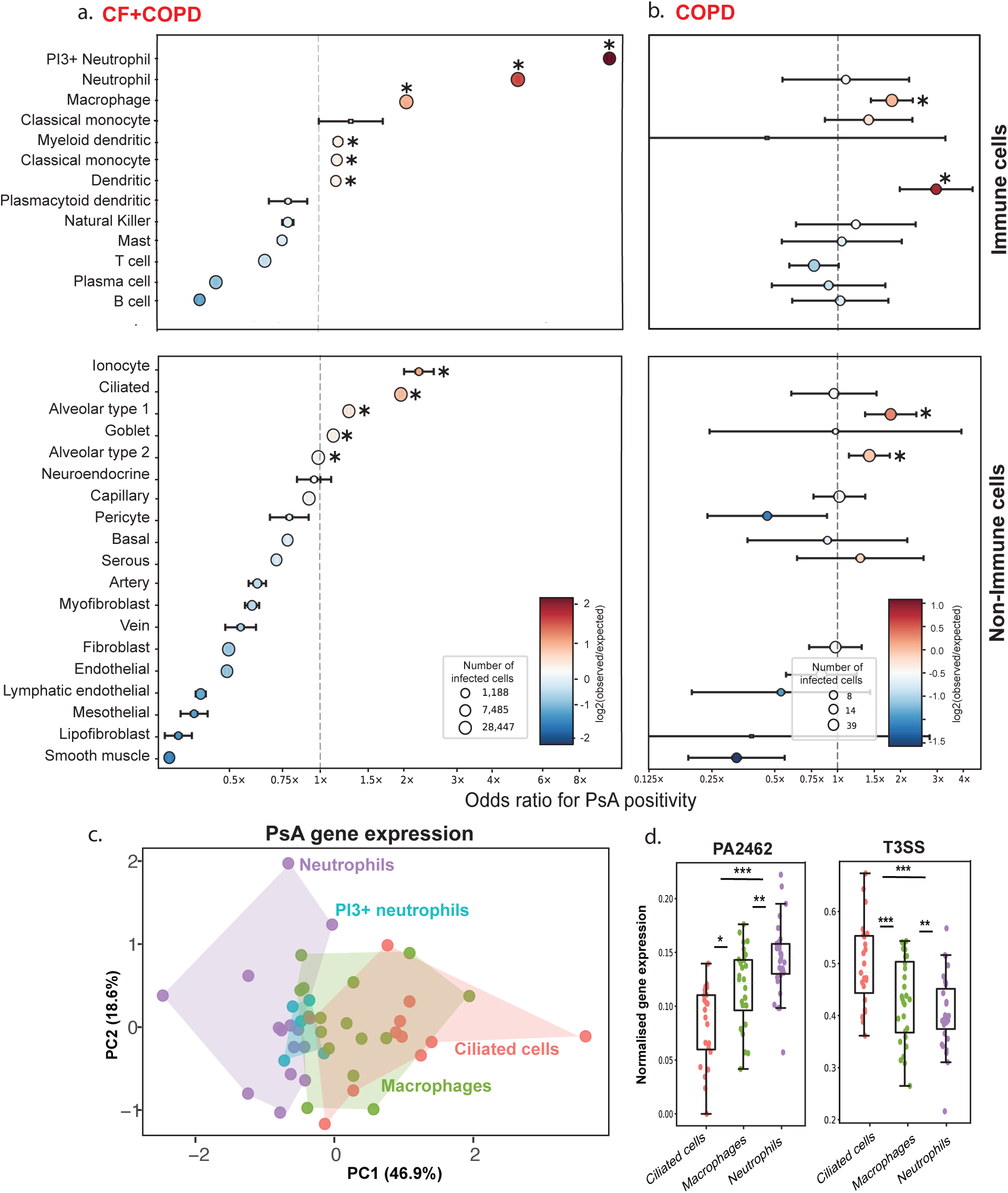
Cell association and divergent gene expression of *P. aeruginosa*. Cells identified in the Xenium datasets were assigned to a cell type using CellAssign with a modified marker list curated through literature searches. PsA = *P. aeruginosa.* a) All cells identified across CF and COPD donors. Cell types with total counts less than 1000 cells were excluded. Cells positive for *P. aeruginosa* were those that had 2 or more transcripts within the cell boundary. A Mantel-Haenszel pooled odds ratios with confidence intervals was calculated across patients. Cell types significantly enriched for *P. aeruginosa* are indicated with an asterisk (adjusted p-value =< 0.05) b) The same analysis was performed for COPD donors only. c) *P. aeruginosa* transcripts detected within phagocytic cells or ciliated cells were extracted and relative abundance was calculated and plotted on a PCA controlling for donor ID. Each point represents the composite profile identified from cells with at least 100 transcripts in a tissue section. d) We performed differential gene expression analysis between the cell types with the total dataset in addition to separate analyses for airway group 1 and airway group 2 donors to account for different *P. aeruginosa* transcriptional profiles described in Fig 4a. Two genes (PA2462 and *exoT/exoS/pcrV* (T3SS)) were significantly different between cell types across all analyses and are plotted. *PI3+* and *PI3-* neutrophils were combined for this analysis.

Neutrophils were strongly enriched for *P. aeruginosa* and were at least fourfold more likely to be bacterial-positive than other cell populations. In particular, *PI3*+ neutrophils showed the greatest enrichment for bacterial transcripts. Other phagocytic populations, including macrophages and monocytes, were also significantly enriched. Among non-immune populations, ionocytes were enriched for *P. aeruginosa* signal; however, most ionocyte-marker-positive cells were dispersed throughout sputum plugs rather than localised to airway epithelium (Extended data Fig. 4), suggesting these represented damaged or fragmented cells. As previously shown in vitro, ciliated epithelial cells were also associated with *P. aeruginosa* signal^15^.

Because CF samples accounted for the majority of detected *P. aeruginosa* transcripts, we performed a separate analysis of COPD donors to determine whether bacterial cell associations differed by disease context (Fig. 5b). Despite substantially lower numbers of bacterial-positive cells, macrophages, dendritic cells, and alveolar type I and type II cells were significantly enriched for *P. aeruginosa* transcripts.

We next examined whether *P. aeruginosa* transcriptional state varied by host cell association. Analyses focused on the four cell populations containing the highest numbers of bacterial-positive cells. *P. aeruginosa* transcript counts were aggregated by tissue section and normalised to total bacterial mRNA abundance. Principal component analysis identified a transcriptional gradient separating neutrophil-associated and ciliated-cell-associated bacterial profiles (Fig. 5c). This separation was driven by differential expression in virulence-associated programmes, including increased expression of *PA2462*, encoding a putative haemolysin and type III secretion system genes (Fig. 5d).

### Elafin (*PI3*)-positive neutrophils are enriched for *P. aeruginosa* across airway and vascular niches

*PI3* (elafin)-positive neutrophils showed the strongest enrichment for *P. aeruginosa* transcripts (Fig. 5a), and this enrichment was observed in both airway and blood vessel compartments in CF tissue (Extended data Fig. 5a). *PI3* expression was restricted to a distinct neutrophil subpopulation rather than broadly expressed across all neutrophils (Fig. 6a) Although *PI3* is classically associated with epithelial mucosal cells, previous studies have also identified *PI3*-expressing neutrophil populations in disease^27,28,29^. Consistent with this, analysis of the Visium data identified strong co-localisation of *PI3* with *CXCL8* (*IL8*) and additional neutrophil inflammatory markers (Fig.6b), potentially reflecting NF-kB dependent regulation of these genes as proposed previously^30^.

**Figure 6.**
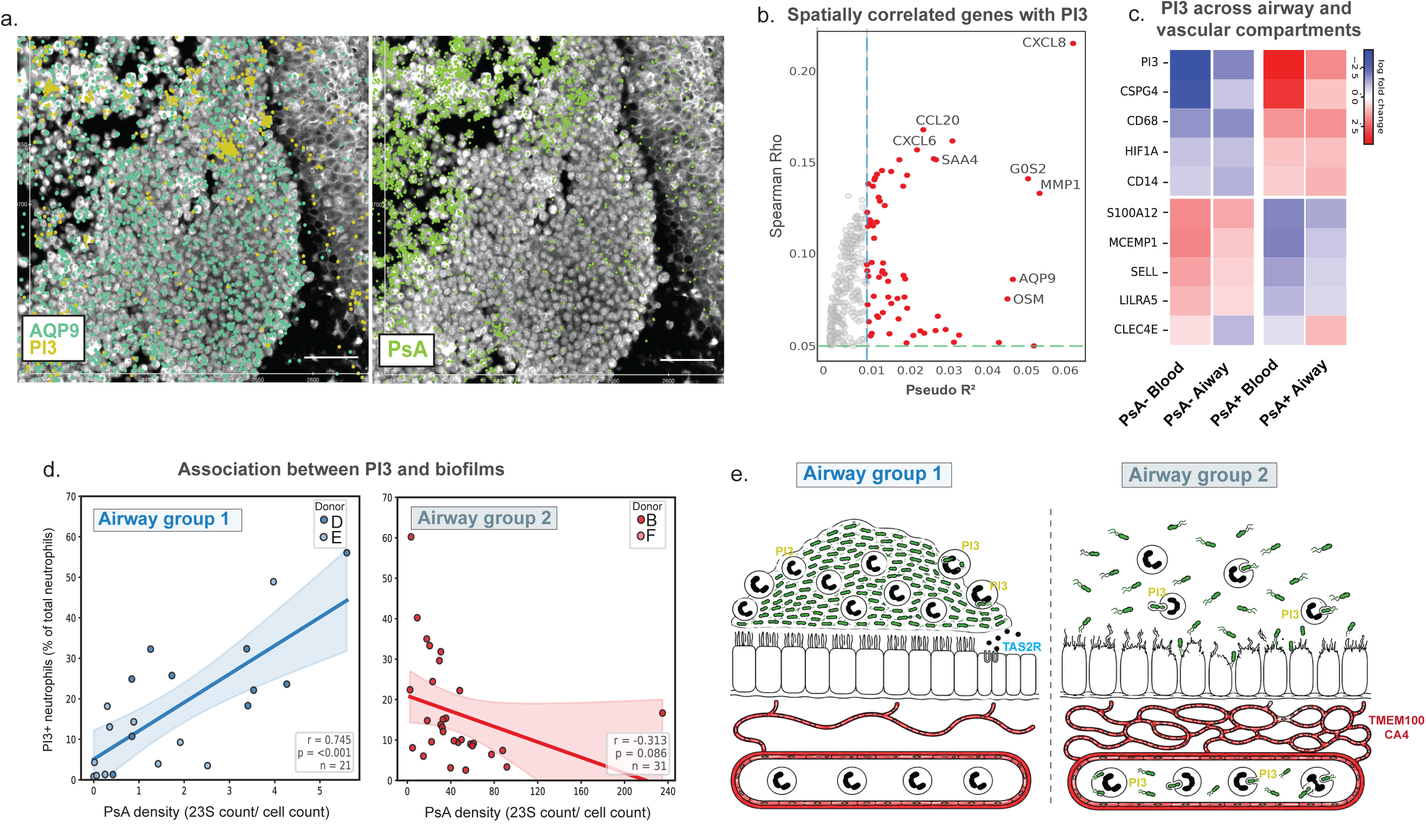
*PI3+* neutrophils are tightly linked with *P. aeruginosa*. PsA = *P. aeruginosa*. a) Images from Xenium Explorer vs4 of a sputum filled airway, showing detection of AQP9 (marker of activated neutrophil) and *PI3* as clustered subpopulations of cells (left) and co-localisation with *P. aeruginosa signal* (right). The scale bar is 50μm. b) Spatial correlation analysis with *PI3* using Visium data. Genes whose expression increases with PI3 are identified by a negative binomial GAM (pseudo R2 ≥ 0.01) and supported by a Spearman correlation (ρ ≥ 0.05), with the top associated (spot-based informed) genes highlighted c) Differential expression analysis comparing *P. aeruginosa* positive and negative neutrophils. Only genes identified as significantly different across both compartments are shown (p < 0.05) *CLEC4E* is the only gene to show a divergent expression profile between the compartments) d) Linear regression analysis between *PI3+* neutrophil proportion and *P. aeruginosa* density in airways using the Xenium data. Density is calculated by dividing *P. aeruginosa* 23s rRNA count by the total neutrophil count to account for different sizes of sputum plug. Each point represents all the annotated airways in a single section. There was a positive significant relationship for donors with predominantly a biofilm-like airway type 1 but not planktonic-like airway type 2. The shaded areas represent 95% confidence intervals. e) Proposed model of airway host–pathogen states in chronic *P. aeruginosa* infection. Schematic showing two spatially distinct airway states identified in chronically infected lung tissue. Left, an alginate-rich biofilm-like state in which dense *P. aeruginosa* aggregates are associated with *PI3+* neutrophil accumulation, limited phagocytosis and host chemical-sensing programmes at the mucosal surface. Right, a motile, active-growth state in which dispersed *P. aeruginosa* interacts more directly with damaged ciliated epithelium and is associated with epithelial stress, inflammatory remodelling, hypervascularisation and intravascular bacteria co-localised with neutrophils.

*PI3* co-localisation with *P. aeruginosa* was associated with an antimicrobial gene module (Extended Data Fig. 5b) and bacterial-positive PI3+ cells resembled canonical activated neutrophils (Extended Data Fig. 5c). Across both airway and vascular compartments, *P. aeruginosa*-positive neutrophils showed increased *PI3* and reduced alarmin-associated genes, including *S100A12* and *MCEMP1,* supporting a model in which bacterial association, rather than anatomical microenvironment, drives the *PI3+* neutrophil state (Fig. 6c).

Given the enrichment of *PI3+* neutrophils around *P. aeruginosa*, we initially expected *PI3+* neutrophil burden to scale with total bacterial load across tissue sections. However, this relationship was not observed (Extended Data Fig. 5d), indicating that bacterial abundance alone does not explain expansion of the *PI3+* neutrophil state. Instead, *PI3+* neutrophil burden scaled with *P. aeruginosa* load only in sections containing airway group 1 biofilm-associated bacteria, but not in sections containing airway group 2 planktonic-like bacteria (Fig. 6d). This suggests that this neutrophil state is preferentially linked to the biofilm-associated *P. aeruginosa* programme. Thus, the *PI3+* neutrophil programme appears to reflect bacterial state-dependent immune reprogramming rather than nonspecific inflammation.

## Discussion

We have found that chronic *P. aeruginosa* infection is spatially structured in human lung tissue, with anatomical and cellular niches linked to distinct bacterial transcriptional states and immune interactions. Concordant with previous histopathological studies^31^, we found that the dominant niche was large airways. Within the airways we noted the presence of two distinct *P. aeruginosa* transcriptional programmes (Fig. 6e), which was not explainable by underlying strain genetics. This is in comparison to previous work that suggested that genetically diverse populations have similar transcriptomic profiles in the CF lung.^9^ Our data suggests that patient, treatment or host factors can modulate the airway niche leading to divergent bacterial lifestyles, rather than a single uniform biofilm state as proposed previously^32–34^.

The *P. aeruginosa* signal from airway group 1 had higher expression of *algE*, responsible for alginate production and upregulated during biofilm production^21,20^, and consistent with this was phagocytosed less^22^ and spatially co-located with higher numbers of neutrophils^23,24^ . This was associated with significant upregulation of sensory perception gene programmes in these airways. This included airway chemosensory receptors known to detect bacterial products, including *P. aeruginosa* quorum-sensing molecules^25^, which likely accumulate at high local concentrations within or around persistent biofilms. Among the upregulated genes was *TAS2R38*, which encodes a bitter taste receptor expressed in airway epithelium. Allelic variation in TAS2R38 has been linked to susceptibility to chronic rhinosinusitis^35^ and *P. aeruginosa* infection^36^ in people with CF and upper respiratory infections^37^. Higher TAS2R38 activity is considered protective because it can enhance epithelial antimicrobial responses^37^. Thus, increased expression of *TAS2R38* and related sensory-response genes reflects a more active epithelial chemosensory defence program in these airways, consistent with host detection of biofilm-associated bacterial molecules.

In contrast, the *P. aeruginosa* signal from airway group 2 was consistent with increased motility, active growth, and higher T6SS activity. These airways had significantly stronger signatures of ciliary stress, inflammation, and epithelial remodelling. Compared with biofilm-associated bacteria, motile and actively growing *P. aeruginosa* are likely to have greater direct interaction with the mucosal surface, while T6SS activity may further enhance host-cell contact and invasion^13^.

Together, these bacterial features contribute to a more inflamed and injury-prone epithelial state. In addition, group 2 airways showed significant enrichment of proteolysis-related pathways, consistent with epithelial damage and increased release or leakage of host proteins into the airway lumen. This protein-rich inflammatory environment may provide additional nutrients that support *P. aeruginosa* growth, reinforcing an acute, actively replicating bacterial state. Notably, donors with this airway state also showed higher intravascular bacterial burden and stronger hypervascularisation signatures, suggesting a link between mucosal injury, vascular remodelling and local access of *P. aeruginosa* to the vascular compartment.

Such a state provides a plausible tissue-level mechanism for pulmonary exacerbation: local transition from a relatively contained biofilm-associated state to a motile, actively growing and epithelial-damaging state could amplify inflammation, impair mucociliary clearance and increase bacterial activity without requiring acquisition of a new strain.

This distinction provides a mechanistic explanation for why genetically related bacteria within the same lung can generate divergent host responses^9,38^. Rather than reflecting strain identity alone, bacterial behaviour appears to be locally conditioned by the airway microenvironment, with alginate-rich biofilm-like states associated with neutrophil-rich inflammatory foci and motile, T6SS-associated states linked to epithelial stress and remodelling.

We found that *P. aeruginosa* transcripts were spatially co-localised with multiple host cell populations, consistent with either intracellular colonisation or attachment to host cells. This association was predominantly neutrophilic in CF tissue, whereas macrophages and dendritic cells were more prominent in COPD, demonstrating disease-specific patterns of host–pathogen interaction. Beyond phagocytic immune cells, *P. aeruginosa* signal was also associated with ciliated epithelial cells, goblet cells, and alveolar type I and type II cells, expanding the range of cellular niches in which *P. aeruginosa* may persist in the human lung.

We found that bacterial transcriptional profiles differed according to host cell association. T3SS genes were more highly expressed in bacteria interacting with ciliated cells, consistent with recent in vitro models proposing that *P. aeruginosa* deploys the T3SS during invasion of the mucosal barrier^13,39^. In contrast, *PA2462* had significantly higher expression in phagocytic cells than ciliated. PA2462 remains poorly characterised, but has been identified as a putative haemolysin^40^ and to contact-dependent inhibition^41^, suggesting a potential role in bacterial competition, adhesion, or interaction with host cells. Our data demonstrates that host-cell context is not simply a passive site of bacterial localisation, but an important determinant of *P. aeruginosa* transcriptional state during chronic infection.

We detected intravascular *P. aeruginosa* signal in all 5 CF donors, expanding the ecological range of chronic *P. aeruginosa* infection beyond the airway lumen. Only a single case of cryptic bacteraemia in CF has been identified through histopathological analysis previously^42^. The donor with the highest intravascular burden in our study had a history of haemoptysis, which has been associated with chronic *P. aeruginosa* infection previously^43^. We find that intravascular colonisation was highly correlated with signals of hypervascularisation^44^ and remodelling in CF, which leads to frequent bleeding and fibrin clots. We propose that vascular remodelling, haemorrhage and fibrin deposition provides protected intravascular scaffolds in which *P. aeruginosa* can localise and creates transient access points to blood vessels. Whether this represents locally restricted intravascular persistence within diseased lung tissue or unrecognised systemic dissemination cannot be determined from our data. However, systemic infection or suspected sepsis would generally be a contraindication to lung transplantation. Moreover, sepsis is extremely rare in CF^42,45^ despite the high bacterial burden and extensive tissue damage characteristic of advanced disease. Previous work has suggested that intense immune surveillance in CF may limit systemic spread or prevent progression to sepsis^46^ . Consistent with this possibility, intravascular *P. aeruginosa* signal in our data was closely associated with immune cells, suggesting active immune containment within this niche. Recent work showed that CF pulmonary exacerbations are accompanied by blood neutrophil transcriptional and epigenetic changes consistent with systemic sensing of *P. aeruginosa*-associated products, including flagellin^47^. This supports the possibility that intravascular or vascular-proximal bacterial material in CF lung tissue contributes to systemic neutrophil activation during exacerbation.

The enrichment of *PI3+* neutrophils provides a link between bacterial localisation, bacterial lifestyle and local immune regulation. *PI3* encodes elafin/trappin-2, an inhibitor of neutrophil elastase with anti-inflammatory^48^ and anti-microbial^49,50^ activity. *PI3* expression has also been associated with long-lived neutrophil states ^23^, raising the possibility that *PI3+* neutrophils are induced by the chronic sputum environment rather than by direct bacterial interaction. However, we identified only limited evidence of co-expression with anti-apoptotic regulatory programmes, and *PI3+* neutrophils were enriched for *P. aeruginosa* across both airway and vascular compartments. In agreement with this, previous studies have identified *PI3* upregulation in both healthy and CF neutrophils exposed to lipopolysaccharide (LPS)^51^. Thus the *PI3+* state of neutrophils is linked closely to bacterial infection.

*P. aeruginosa*-positive neutrophils displayed increased *PI3* expression alongside genes involved in antimicrobial responses, together with reduced expression of otherwise highly expressed alarmins such as *S100A12*. This pattern is consistent with a shift towards a modified inflammatory state rather than simple neutrophil activation. The stronger association of *PI3+* neutrophils with biofilm-associated bacterial profiles further suggests that neutrophil state may reflect bacterial lifestyle. We have shown that biofilm-associated *P. aeruginosa* recruit large numbers of neutrophils while resisting phagocytic clearance, generating dense neutrophil-rich inflammatory foci and high local protease burden.^52^ In this context, *PI3* expression may represent a host attempt to ameliorate neutrophil elastase-mediated tissue injury as previously proposed in the context of convalescent COVID-19^29^. However, proteomic studies have found that elafin is often cleaved and therefore functionally impaired in the CF airway^53^, which may limit this protective response. These observations suggest that the *PI3*–elafin axis may represent an endogenous but potentially insufficient attempt to limit protease-mediated tissue injury in biofilm-associated infection.

We have advanced spatial transcriptomic technologies to allow us to profile both host and pathogen activity across a physiological environment in a single experiment. Our data support a model in which chronic *P. aeruginosa* infection is maintained by spatially structured bacterial subpopulations that occupy distinct anatomical and cellular niches, adopt different behaviours, and influence immune cell state. Chronic *P. aeruginosa* persistence may therefore arise not from biofilm tolerance alone, but from the ability of bacteria to partition into spatially distinct physiological states that are maintained by local host tissue environments. Therapeutic strategies that treat chronic infection as a single airway biofilm may therefore miss niche-specific bacterial states.

## Author Contributions

J.M.B and R.A.F conceived the study. C.K and J.M.B initiated and led the study. J.M.B, S.A.T and M.P supervised the work. C.K, K.R, L.E, K.T, J.H, H.M.C, V.B.P, B.R, H.A, S.U, M.F and T.K.J performed experiments and generated data. N.J.C performed imaging and histology. C.K, D.P, P.B, A.B, S.D and J.M.B performed formal analysis. H.W contributed histopathological interpretation. S.A.T and O.B contributed spatial analysis methodology and interpretation. R.A.F contributed clinical and biological interpretation. J.M.B and C.K wrote the original draft. All authors reviewed and edited the manuscript. J.M.B and O.B acquired funding.

## Acknowledgements

We thank the donors, their families and the staff at the Royal Papworth Tissue Bank. This research was funded in whole, or in part, by the Wellcome Trust 220540/Z/20/A. Muhanad Tuameh (Wellcome Sanger Institute) provided useful input for the bacterial clustering methods we implemented. We thank Professor Stephen Bentley for his useful feedback on the manuscript. We wish to thank the core facilities at the Wellcome Sanger Institute (DNA pipelines, spatial core facility, Parasites and Microbes programme core lab, informatics and samples management) and Martin Prete for their contributions to this work. For the purpose of Open Access, the author has applied a CC BY public copyright licence to any Author Accepted Manuscript version arising from this submission.

S.A.T. is a scientific advisory board member of Bioptimus, ForeSite Labs, Xaira Therapeutics, a co-founder, Board observer and equity holder of TransitionBio, a co-founder, consultant and Board Director of Ensocell Therapeutics, a non-executive director of 10x Genomics and a part-time employee of GlaxoSmithKline.

## Data availability statement

All source data for figures is provided in a supplementary document. Sequencing data has been deposited under the following accessions: EGAS00001008178 and ERP149661.

