## Extended data figures for "Spatial lung niches shape *Pseudomonas aeruginosa* persistence"

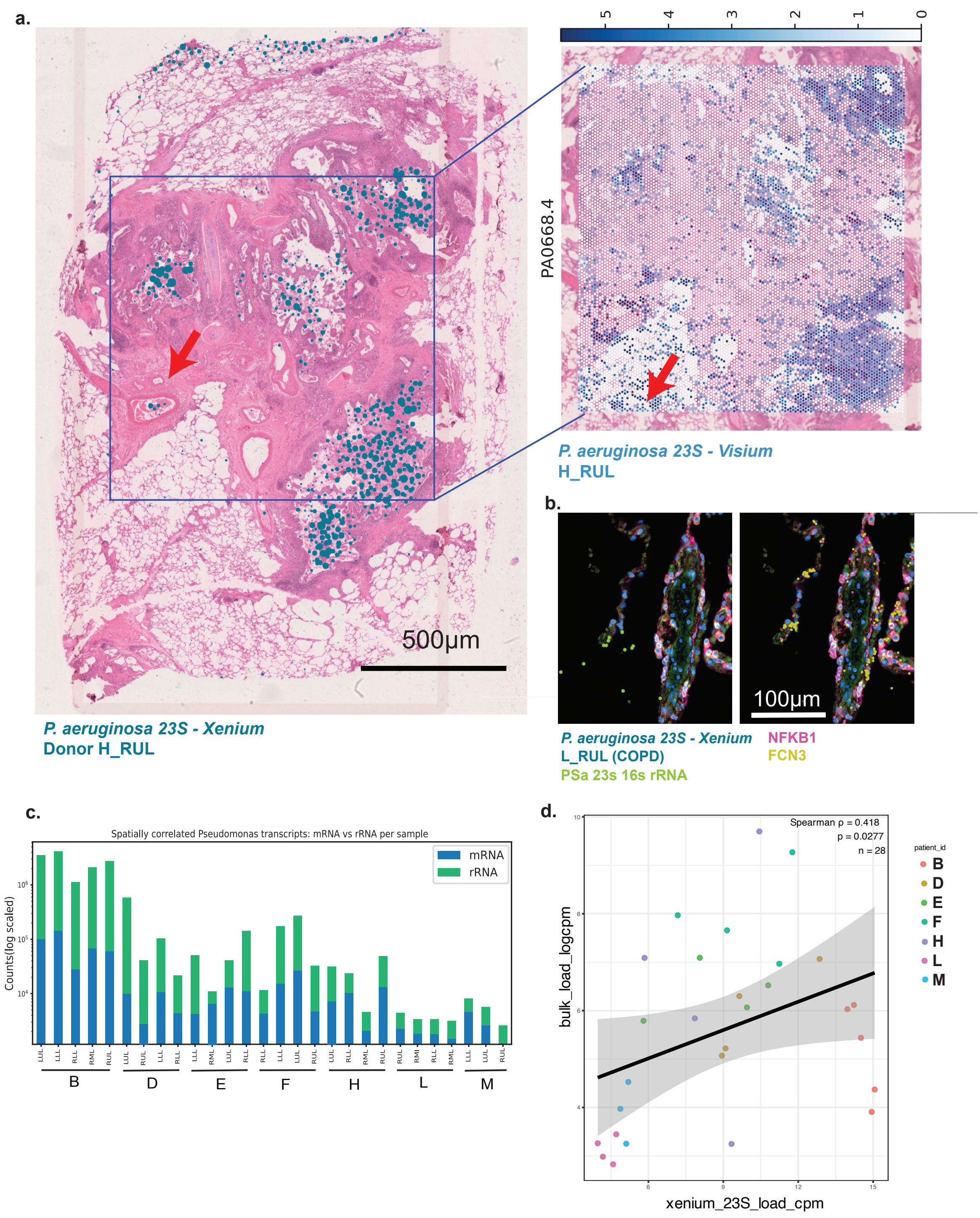


**Extended Figure 1.** Application of host-pathogen spatial transcriptomics. a) Whole slide H&E image of Donor_H_RUL with *P. aeruginosa* signal overlayed (blue) demonstrating localisation to the major airways. Highlighted area is displaying Visium with Cytassist data on region of interest from the same section shown in demonstrating concordant signal between the two technologies. b) Example Xenium image from a COPD lung section showing co-localisation of *P. aeruginosa* transcripts and immune markers in the alveolar parenchyma. c) Counts of spatially co-located P. aeruginosa transcripts across donors and tissue sections. d) Linear regression analysis between Xenium *P. aeruginosa* 23S and bulk *P. aeruginosa* load in the RNAseq data (counts per million). 23s rRNA is used as a measure for *P. aeruginosa* load throughout because in one donor the 16s probe appeared to give poor signal.

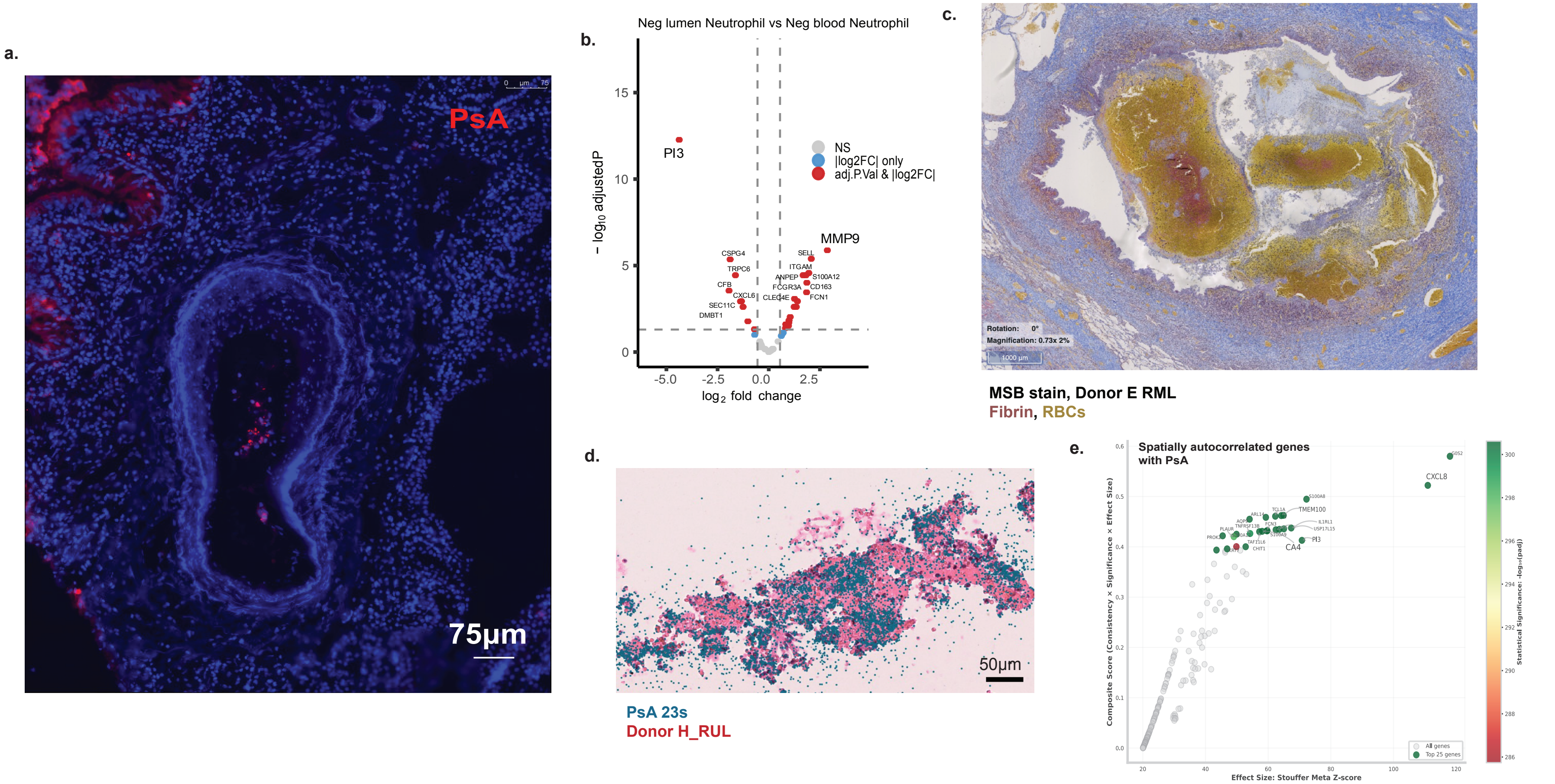


**Extended Figure 2.** Exploration of vascular *P. aeruginosa* (PsA) signal. a) Immunofluorescent image of Donor B RUL blood vessel using a anti-*P. aeruginosa* polyclonal primary antibody. b) Differential gene expression analysis for PsA negative neutrophils between airway and vascular niches, showing niche specific specialisation. c). Martius, Scarlet, and Blue (MSB) strain demonstrating high deposition of blood and fibrin in sputum plugs. d) Overlap of Xenium PsA signal onto fibrin deposit (H&E) found in an airway. e) Spatial autocorrelation analysis on Visium data with PsA identifying significant co-localisation with immune signals and genes associated with vascularisation (TMEM100 and CA4) rating localisation to the major airways.

**Extended Figure 3.**

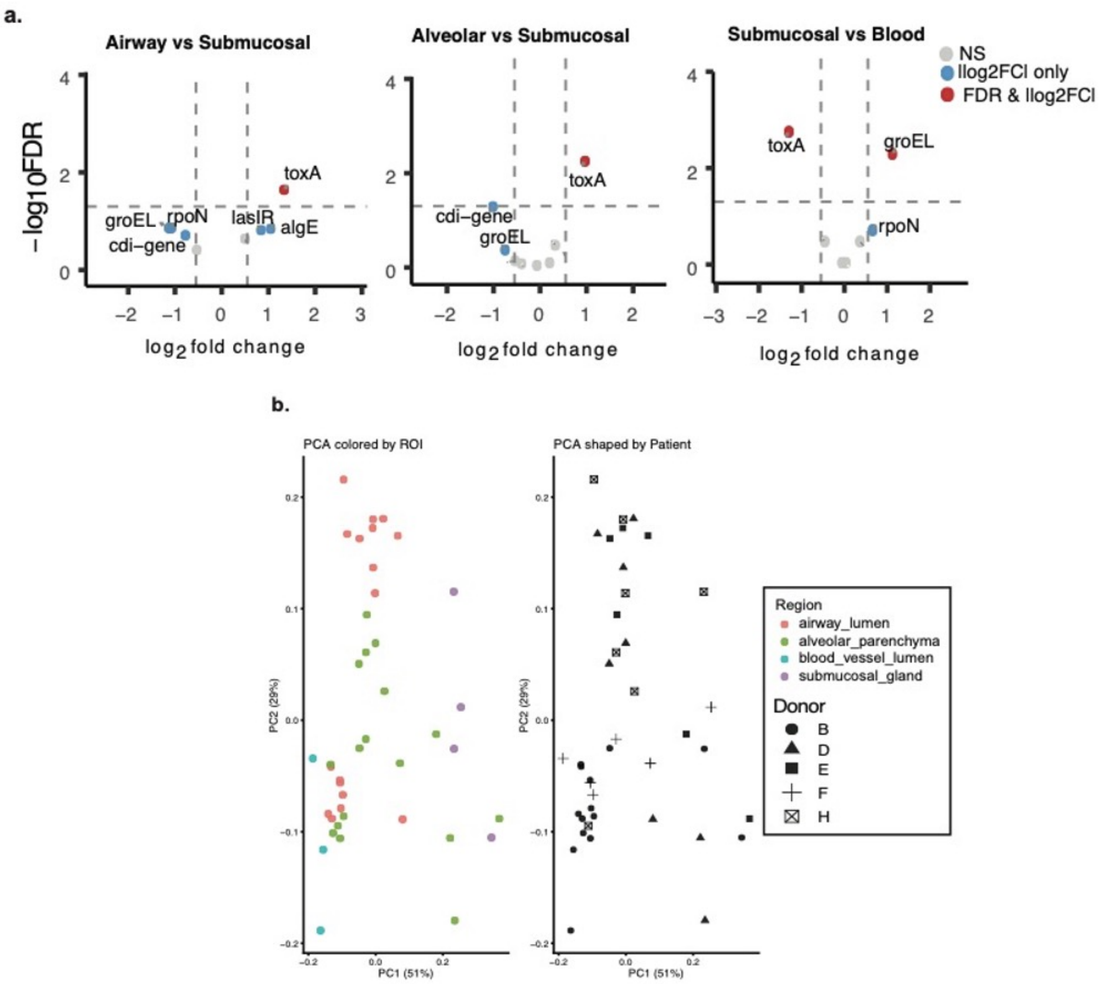


Differential expression analysis between anatomical niches. A) Volcano plots showing differential *P. aeruginosa* gene expression in anatomical niches. Red dots indicate genes that had greater than 2-fold change and were statistically significant. b) PCA analysis shown in Figure 4a but labelled by location and donor ID

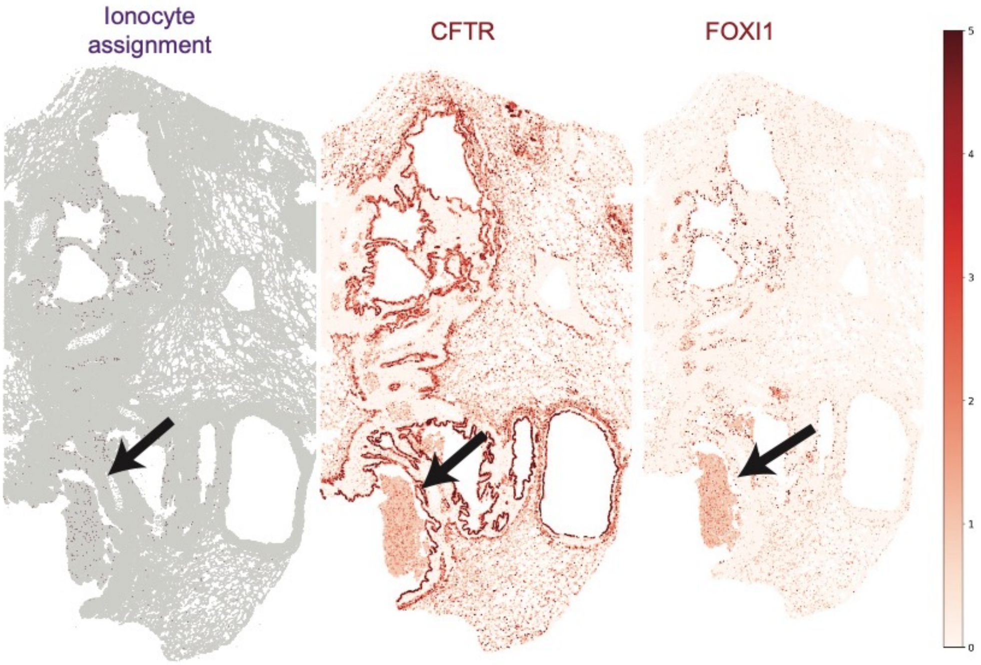


**Extended Figure 4.** Example Xenium section from CF donor showing cells assigned as ionocytes based on expression of canonical markers CFTR and FOXI1. Ionocyte-assigned cells were predominantly detected within dense sputum-rich regions (black arrow) rather than along the epithelial mucosal surface, where ionocytes would be expected anatomically. This distribution suggests that ionocyte assignments in these regions likely reflect epithelial cell fragmentation and/or ambient epithelial RNA within inflamed sputum, rather than intact ionocytes.

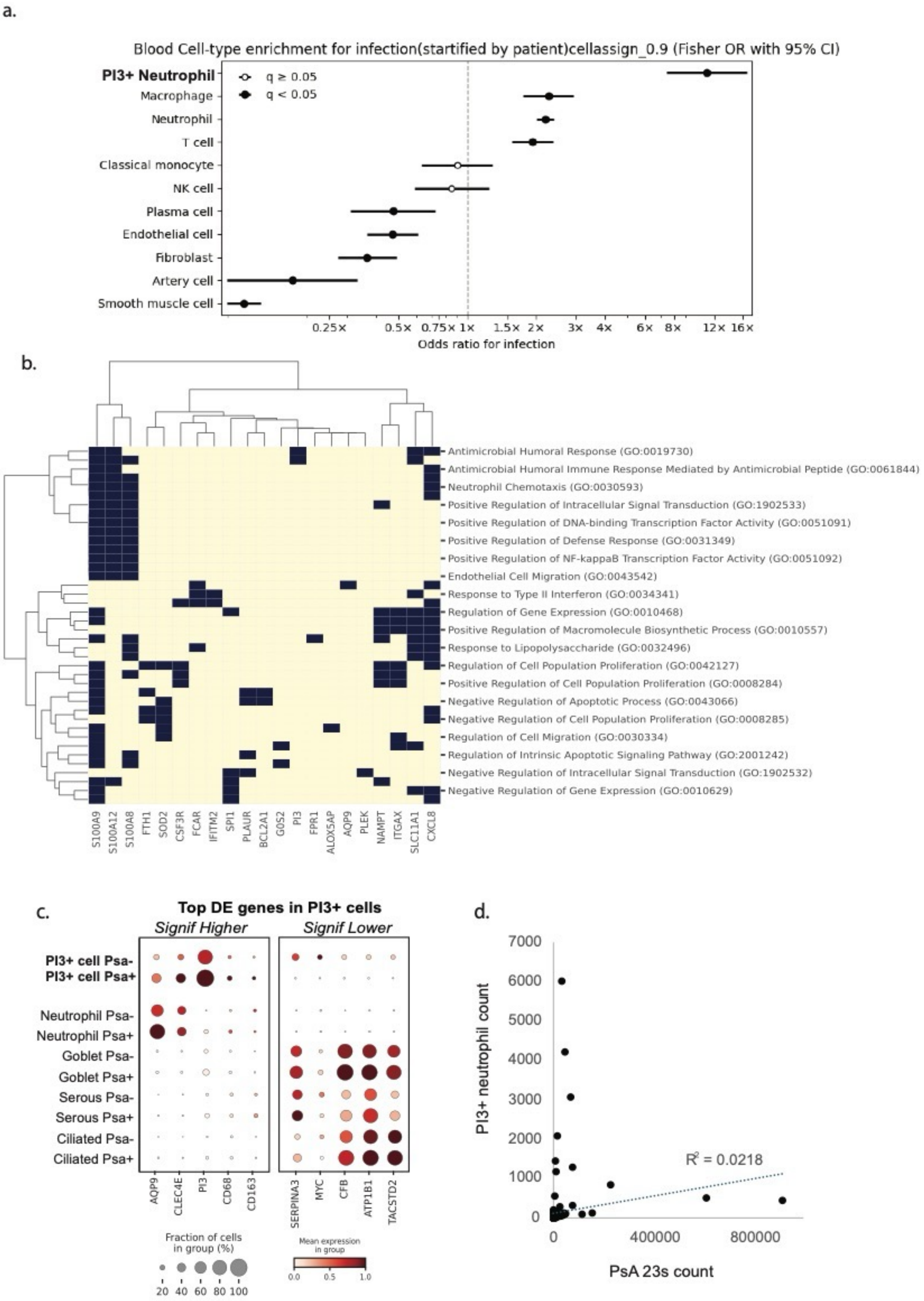


**Extended Figure 5.** Relationship between PI3+ neutrophils and *P. aeruginosa*. a) Cell association analysis for blood vessel lumen regions showing similar enrichment for PI3+ neutrophils. Cells positive for *P. aeruginosa* were those that had 2 or more transcripts within the cell boundary. A Mantel-Haenszel pooled odds ratios with confidence intervals was calculated across patients. Cell types significantly enriched for *P. aeruginosa* are indicated with an asterisk (adjusted p-value =< 0.05). b) Spatial autocorrelation analysis was performed to identify genes that were spatially collocated with *P. aeruginosa* transcripts in the Visium data. Spatially autocorrelated genes were first identified using a depth-adjusted negative binomial model and retained at FDR < 0.01. Pairwise local correlations between retained genes were clustered to define spatial gene modules, requiring at least 10 genes per module. Module activity was scored across spatial locations, and top-loading genes were functionally annotated using EnrichR. Shown here is a presence/absence map of genes and their predicted functions in the module containing PI3, which is enriched for genes associated with antimicrobial function. c) Differential gene expression analysis showing the top five highest and lowest expressed genes in PI3+ *P. aeruginosa*+ cells compared to PI3+ *P. aeruginosa*- cells. All were significant with adjusted P values < 0.001. Relative expression is shown for other cell types to demonstrate that PI3+ *P. aeruginosa*+ cells resemble canonical neutrophils. d) Linear regression analysis between PI3+ neutrophil and PsA load. The correlation was non-significant P>0.05.
