## Supplementary methods for "Spatial lung niches shape *Pseudomonas aeruginosa* persistence"

**Donor Tissue**

Human lung tissue was obtained from the Royal Papworth Tissue Bank under approved tissue-bank governance (REC approval 23/EE/0198). The tissue bank obtains consent for research use of donated tissue, and samples were provided to the study in de-identified/pseudonymised form. Formalin Fixed Paraffin Embedded (FFPE) and frozen lung blocks were requested for donors with COPD or CF who had a known clinical history of *P. aeruginosa* chronic infection and the associated metadata is found in Supplementary Table 1.

**Bulk RNA extraction and sequencing**

Bulk RNA was extracted from 3 x10 micron FFPE lung scrolls using RNeasy (Qiagen) extraction kit, per manufacture recommendations. Distribution of RNA fragment sizes (DV 200) were determined using the HighSensitivity RNA ScreenTape (Agilent). Both 18S (human) and 16S (bacterial) were selectively removed via probe hybridisation using the NEBNext® rRNA Depletion Kit v2 (NewEngland Biolabs) and NEBNext® rRNA Depletion Kit (Bacteria) respectively. We reasoned that since the RNA was quite fragmented and poly-A tails might be loose, we omitted poly-A depletion and additional fragmentation steps. Paired-end Illumina libraries were prepared and sequenced 12-plex on Novaseq S4 XP flow cell.

**Bulk human cell composition inference**

Cell type abundances were inferred from processed bulk RNA-seq data using CIBERSORTx [1] with batch correction and quantile normalisation disabled, relative run mode, and 1000 permutations. Non-immune cell type composition was determined using the extended Human Lung Cell Atlas (level 3 annotation) reference [2]. Immune cell type composition was determined using the LM22 reference [1].

**Metatranscriptomic analysis**

Human reads were aligned using STAR version 2.7.10b against Homo_sapiens.GRCh38.112. The remaining “non-human” reads were then trimmed by using Trimgalore version 0.4.4 [3]. The resulted trimmed reads were taxonomically profiled, by running Kraken2 and Bracken [4] against the standard 16GB kraken2 database (which contains genomes of human, viruses, and bacteria), with the following settings: k = 50, threshold = 100.

We identified contaminants in the samples through multiple approaches. Principal component analysis (PCA), with taxon relative abundance as input, was initially performed to explore variation in the dataset and showed that the largest sources of variance were associated with batch effects. Fourty-seven species with the highest loadings (<-0.1 or >0.1) correlated with batches and were therefore added to a blacklist to minimise batch-associated variation suggestive of sample preparation contamination. In addition we employed Sylph [5] which is a marker based classification tool and therefore used to identify DNA contaminants where there is broad coverage across the genome, which led to the removal of an additional eight species.

To extract gene expression information from the *P. aeruginosa* transcripts in the bulk RNAseq data we first mapped it against the PAO1 genome using BWA-mem [6]. To ensure the reads were *P. aeruginosa* and not close orthologs in other bacterial species we then classified these reads using kraken2 [4] and only used reads that were assigned to *P. aeruginosa*. The number of reads mapping to each gene was counted. And any genes with more than two independent reads (or one read pair) were classified using COG and PseudoCAP [7].

**Bulk DNA extraction, sequencing and bait capture**

Total genomic DNA was extracted from 3 x10 micron FFPE lung scrolls using Qiagen DNA FFPE Advanced kit with UNG (uracil-N-glycosylase).

For targeted enrichment of *P. aeruginosa* DNA we used Agilent Sureselect DNA with a custom bait design. The bait panel comprised of 192,512 120nt probes which tile across the 6.3 Mb P. aeruginosa genome. The baits were designed using PanArray [8] based on three *P. aeruginosa* complete reference genomes (PAO1, PAK and PA14) selected for their completeness and genetic diversity across the species. The resultant libraries were sequenced on a Novaseq S4 XP lane (34-plex) with paired reads of 150bp.

Raw reads were mapped to using BWA-MEM [6] to the PAO1 reference genome and variants called using an in-house pipeline (https://github.com/sanger-pathogens/strain_mapper/tree/v1.7.1). In order to place them in phylogenetic context we selected 25 representative strains from clones common to chronic lung disease as described in a previous publication [9] which were also mapped to the PAO1 reference genome using the same approach. A phylogeny of variable positions for each alignment reconstructed using RAxML (v8.2.8) [10] with 100 bootstraps. Pairwise SNP distances were estimated using SNP-dists (v0.7.0) (<https://github.com/tseemann/snp-dists>).

**Xenium custom probe design**

For Xenium custom padlock probes, probes against target sequences were designed in partnership with 10X using the internal 10X Xenium Panel Designer*.* Candidate probes were checked for specificity to *P. aeruginosa* using NCBI BLASTn [11, 12] with stringent filtering. A probe was considered a suitable candidate if it matched *P. aeruginosa* specifically and produced fewer than 10 off-target hits in other species. Using these criteria, we retained 136 candidate probes across 13 genes, with multiple probes represented for each target. For genes with more than 20 suitable probes, we retained only the 20 probes with the fewest off-target hits.

**Visium custom probe design**

For Visium, we designed probes against similar targets as done in Xenium. Visium probes were designed using an in-house probe design pipeline (https://github.com/sanger-pathogens/lung-microbiome-visium) in line with 10X guidelines for Visium probe design detailed below.

Required inputs for the probe pipeline are; gene file in fasta format containing coding sequences for pre-selected genes for the Visium probe panel; a reference genome in fasta format for the target organism; a file showing coordinates of all SNPs known to be present in at least 5% of genomes in this organism’s population relative to the reference genome.

Firstly, we generated all possible 50-bp sequences, hereafter referred to as probes, using a sliding window spanning the entire range of the target gene. These probes were filtered to Visium requirements, the central base must be a thymine (T) and the two halves of the probe (bases 1-24 and 26-50) must have a G/C content within 44-72%. The passing probes were checked against the input reference and all matching coordinates identified. Probes identified outside of the target gene region were excluded otherwise, the probe was split into three sections, a central zone falling between 22-29bp and a left and right wing either side. Using the user-provided SNP coordinates, all probes with SNPs within the central zone were excluded. In addition, any probes with more than 5 SNPs in either wing also failed. Passing probes were ranked by SNP count and those with the fewest SNPs were prioritised downstream.

The resulting probes from these filters were considered our final probe candidates. These probes were compared first against GTDB 214 genome representatives using pyjellyfish (<https://github.com/iric-soft/pyJellyfish>) 1.3.1 [13] any identical matches outside of the target species were discarded. As a final check probes were aligned using blast 2.15.0 [11] against the core_nt (v4) database and any probes with matches against off-target organisms were discarded. The final panel of probes to be used for Visium probe manufacturing were picked from the passing candidate probes based on positional information to maximise coverage over the target gene and the SNP positions to minimise the SNP count occurring within the probe. We retained 29, 50bp, suitable probes against 11 target genes RHS and LHS probes were modified as per 10X guidelines and probes ordered from IDT as o-pools.

**Xenium processing – Human Lung panel v1 (289 genes) with custom probe spike-in**

FFPE blocks were sectioned to thickness of 5um onto Xenium slides. Sections were centrally placed within the sample area (10.45 mm x 22.45mm), air dried and baked to prevent tissue detachment. This was followed by a series of xylene and ethanol washes to deparaffinise the sections. Sections were decrosslinked to ensure target retrieval. Resuspended custom add-on probes (10 ul) were mixed with Xenium pre-designed gene expression panel and probe hybridisation buffer to make a final hybridisation mix. The probe hybridisation mix was applied to processed tissue sections and incubated overnight. The following day, unbound probes washed off using a series of washes. Bound probes were ligated followed by rolling circle amplification. Sections were incubated overnight at 4C with the Xenium Multi-Tissue Stain Mix, a four-channel cocktail of antibodies for cell boundaries and cytoplasmic: epithelial cell membrane (ATP1A1 and E-Cadherin), CD45 for immune cell boundaries, 18S ribosomal RNA probe for cytoplasmic identification and alphaSMA and Vimentin for stroma and connective tissue. Following staining and autofluorescence quenching, nuclear membranes were counterstained with DAPI.

Xenium Onboard Analysis (Software version 3.2.1.2) was used for multimodal segmentation. For downstream analysis we considered only transcripts with high quality (Phred quality score >= 20).

**Xenium downstream analysis**

Xenium spatial transcriptomics downstream analysis was performed using standard SpatialData (0.5.0) [14] and Scanpy (1.11.5) [15] workflows. Due to a low median number of transcripts per cell (across 28 samples median transcripts ranged from 12 to 67 transcripts/cell), consistent with fibrosis from end-stage disease, and low expected transcript abundance in neutrophils [16], moderately stringent quality filters were applied, retaining cells with *min_transcripts* ≥ 3. Xenium/spatial transcripts best-practice recommendations were incorporated throughout preprocessing, and rapids-singlecell (0.13.1) [17, 18] was used to support analysis at scale (millions of cells). Cells were normalized to a target sum of *1e2* and log1p-transformed, followed by PCA (30 components). Neighbourhood graph construction used *n_neighbors = 15*. UMAP embeddings were generated with *min_dist = 0.1*.

Cell annotation was performed using CellAssign [19] owing to a limited set of human gene targets on the panel. Only cells with high-confidence assignments were retained (confidence > 0.9) for downstream analyses. Neutrophils were defined using canonical markers (ITGAM/CD11B, AQP9, S100A12 [20, 21], as this population was absent from the human lung v1 panel and not represented in the human lung atlas [22]. Rare ionocytes were confirmed by FOXI1 expression. Clusters labelled as monocytes were renamed to “unassigned/ambiguous” when marker expression was inconsistent and proliferative labels were reprofiled into cell types after masking proliferation-associated marker genes

**Spatial correlation**

Spatial interactions between *16S* and *23S* rRNA transcripts were assessed using the spatstat (3.4-0) [23]. For each sample, transcript coordinates were imported and filtered to retain only *16S* and *23S* transcripts of *Pseudomonas aeruginosa*. To control computational load, datasets exceeding 100,000 points were randomly subsampled. Spatial point patterns were constructed with transcript coordinates defined point locations and transcript type (*16S* or *23S*) was encoded. The observation window was defined as the convex hull enclosing all detected transcript locations.

Bivariate spatial dependence between 16S and 23S transcripts was quantified using Ripley’s cross-K function [23]. Monte Carlo simulations (n = 99) were performed to generate confidence envelopes under the null hypothesis of spatial independence between the two transcript types.The observed K-function was compared against the simulation envelopes to evaluate significant spatial attraction or repulsion across distance scales. Based on this analysis, 16S or 23S transcripts lacking a neighboring counterpart within 20 µm (~100 pixels) were considered likely technical noise. We therefore applied neighborhood-based filtering to retain only 16S and 23S transcripts located within this radius cutoff.

***P. aeruginosa* cell type enrichment**

We adapted the microbial single cell approach employed by CSI microbe [24]. To determine cell-type enrichment, we used the donor-specific background infection rate to calculate the expected number of infected cells for each cell type, tested whether each cell type was overrepresented among infected cells within each donor, and then combined evidence across donors.

Cells were classified as *P. aeruginosa* transcript-associated if their segmentation boundary contained ≥2 filtered P. aeruginosa transcripts. For each cell type, enrichment among bacteria-positive cells was tested in a donor-stratified framework. Within each donor, 2 × 2 contingency tables were constructed comparing the number of infected and uninfected cells of the test cell type against all other cell types, and one-sided Fisher’s exact tests were performed. Cell types comprising <1000 total cells were excluded. Per-donor p-values were combined across donors using weighted Stouffer’s method, with weights based on the expected number of infected cells under the donor-specific background infection rate.

Effect size was defined as the log2 ratio of summed observed to summed expected infected cells across donors, with confidence intervals estimated using a Poisson approximation. A Mantel–Haenszel pooled odds ratio and confidence interval were also calculated across donor strata. Multiple-testing correction was performed using the Benjamini–Hochberg method.

**Differential gene expression analysis**

*P. aeruginosa* differential expression between anatomical regions was assessed using edgeR [25] and limma [26] on paired region-level count data. Region group was set a categorical variable, and a design matrix was specified with pair as blocking covariate and region group as the main effect of interest. Since we had a limited custom panel, all eleven *P. aeruginosa* genes on the custom gene panel were retained for the analysis. Library sizes were computed, and normalization factors were calculated using the TMM (Trimmed Mean of M-values). Differential expression was tested using edgeR’s quasi-likelihood negative binomial generalized linear model framework. Differential expression for submucosal glands was then tested relative to all other regions, P-values were adjusted for multiple testing and genes with significant differential expression identified.

To compare gene expression between *P. aeruginosa* negative and positive neutrophils in blood and lumen compartments, neutrophil-specific differential expression between blood and lumen groups was performed using a paired pseudo bulk voom-dream approach [27, 28]. Neutrophils with >= 2 bacterial transcripts were considered positive/infected. For each sample, neutrophil counts were aggregated by region type into a gene-by-sample count matrix and matched to corresponding metadata. Metadata variables including donor ID, sample ID and compartment were encoded as factors. Anatomical region was modelled as a two-level factor with lumen as the reference level and blood as the comparison group.

Genes with insufficient expression across the compared groups were removed and the filtered count matrix was analysed. Differential expression was modelled with compartment (region category) as the fixed effect of interest while donor ID and sample ID were included as covariates (random effect terms) to account for correlation among observations from the same donor. Gene wise linear mixed model were fitted using dream.

To compare *P. aeruginosa* gene expression in select cell types, likewise a pseudo bulk voom-dream approach was used with gene-by-sample count matrix aggregated at the sample level. Differential expression was modelled with cell type as the fixed effect of interest while donor ID and sample ID were included as covariates (random effect terms) to account for correlation among observations from the same donor.

**Post Xenium - Visium Cytassist**

Post Xenium, selected sections were taken forward to the Visium Cytassist. Briefly, sections were destained in 0.1N HCl and stored in 1X PBS at 4C overnight. Sections were washed with pre-hybridisation wash followed by probe hybridisation (with custom spike-in as per 10X recommendations) and ligation steps. Regions of interest were captured with a capture area 11 mm by 11 mm using the CytAssist. Capture regions selected to include a variety of anatomical features as well as cover areas with high *P. aeruginosa* signal on Xenium. Standard Cytassist workflow was followed downstream. Resultant libraries and submitted for sequencing 10-plex on a S4 flow cell

**Spaceranger mapping for Visium**

Custom reference for spaceranger (version 2.1.0) mapping was prepared as per 10x guidelines using spaceranger mkref and PAO1 reference genome, GTF files and custom probe files.

After spaceranger mapping, 196,382 spots across 14 samples were obtained with a median of 3,481 counts per spot. Downstream analysis was performed using standard Scanpy (1.11.5) [15] workflow including noise cells/genes filtering, normalization, log1p transformation and then performing data integrated using Harmony [29] (0.0.10). For mapping cell type information, we applied cell2location (0.1.4) [30] and using Human Lung Cell Atlas extended version scRNAseq dataset [2, 22] as the reference.

**Gene modules analysis on Visium**

To identify genes exhibiting consistent spatial correlation with the *Pseudomonas aeruginosa 23S* gene across samples, Moran’s R statistics were first computed independently for each Visium sample using a radial basis function (RBF) spatial weight matrix [31]. For each gene, Moran’s R values, corresponding Z-scores, and p-values were calculated relative to the expression of the target gene.

Genes were filtered within each sample based on statistical significance (p-value < 0.05) and positive spatial association (Z-score > 0). Results from all samples were then aggregated, and genes were ranked based on both the consistency of significance across samples and the magnitude of spatial association. Specifically, for each gene we computed (i) the number of samples in which it was significantly correlated, and (ii) a meta Z-score using Stouffer’s method. Genes were prioritised by the number of significant samples followed by the meta Z-score. From this ranked list, the top 179 genes were selected for downstream analysis.

We applied the Hotspot framework [32] to identify spatial gene modules. First, local spatial autocorrelation was computed for each gene using a depth-adjusted negative binomial (DANB) model [33]. Genes with significant spatial autocorrelation (FDR < 0.01) were retained, resulting in 179 genes for pairwise analysis. Next, pairwise local correlations between genes were calculated to capture shared spatial expression patterns.

Gene modules were then defined by clustering the local correlation matrix, with a minimum gene threshold of 10 genes per module. This resulted in multiple spatial gene modules representing coordinated expression programs. Module activity scores were computed for each spatial location by aggregating expression of genes within each module. Finally, gene loadings within each module were used to interpret module composition, and functional enrichment analysis using EnrichR [34] was performed on top-loading genes to assign biological meaning to each spatial gene program using the Gene Ontology Biological Process (GO BP, version 2026) database [35].

**Spatial niche identification and differential gene expression**

We applied CellCharter (0.3.7) [36] to our Visium spatial transcriptomics data to identify and characterize spatial cellular niches across all CF samples. CellCharter uses an integrated framework, including spatial clustering, batch-effect correction, and geometric neighborhood measurements to robustly define niche structures. The analysis resulted in seven spatial niches with k=7 as the most stable number of clusters. We then extracted the sputum niche, enabling us to focus our downstream analysis on bacterial-related transcriptional programs.

We compared the transcriptional profiles of the sputum niche between donor groups D, E, H and B, F. Using Scanpy’s rank gene groups approach, we performed differential expression with a T-test, and retained only genes with p-value < 0.05 and |log2FC| > 3 to define significantly up- and down-regulated targets. We then used EnrichR [34] for pathway enrichment of the up- and down-regulated gene sets against Gene Ontology Biological Process (GO BP, version 2026).

***P. aeruginosa* Immunofluorescence**

Samples were first frozen by immersion in a slurry bath of isopentane and subsequently embedded in optimal cutting temperature (OCT) compound. The embedded tissue blocks were stored at −80 °C until sectioning. Cryosections were then cut at a thickness of 15 µm and mounted onto glass slides, with two serial sections placed per slide (one positioned in the upper half and one in the lower half of each slide).

All sections were initially rinsed three times in phosphate-buffered saline without calcium and magnesium (PBS−/−) supplemented with 0.005% Tween-20 (PBS-T) to remove residual OCT. Tissue fixation was then performed using 4% paraformaldehyde for approximately 5 minutes to inactivate potential biohazards and preserve tissue morphology.

Sections were incubated in PBS-T containing 1% BSA for the same duration. Following incubation, slides were washed twice with PBS-T. Blocking was performed using 5% BSA in PBS-T for 1 hour at room temperature, followed by overnight incubation at 4 °C. For each experimental block, one of the paired serial sections was reserved as a secondary antibody only control and incubated without primary antibody to assess non-specific background staining. The following day, slides were washed three times with PBS-T and incubated with a rabbit anti-Pseudomonas aeruginosa polyclonal primary antibody (1:500; Abcam, ab68538) [37] for 1 hour at room temperature, with an additional incubation period of up to 2 hours at 4 °C to enhance antigen binding. After primary incubation, slides were washed three times in PBS-T.

Detection was performed using an Alexa Fluor 647-conjugated donkey anti-rabbit secondary antibody (Invitrogen) diluted according to manufacturer instructions and applied for 30 minutes at room temperature. Slides were then washed a further three times with PBS-T. Nuclei were counterstained with Hoescht, followed by two final PBS-T washes. Sections were mounted using appropriate aqueous mounting medium and stored protected from light until imaging.

Imaging was carried out using a Leica Thunder system, with identical acquisition settings applied across all samples, including a 438 nm laser for Hoechst visualisation and 575 nm excitation for Pseudomonas detection, with identical acquisition settings applied across all samples to ensure comparability. Tissue autofluorescence, particularly within the blue and green channels, was recorded and utilised as a reference signal for tissue area delineation.

**Martius Scarlet Blue staining**
FFPE tissue blocks were sectioned at 4μm and mounted onto glass slides. Control sections obtained from UK NEQAS Cellular Pathology Technique control material were also sectioned at 4μm and processed in parallel with all test sections. Sections were incubated at 55–60 °C for 10–60 min to soften paraffin wax prior to deparaffinisation in three changes of xylene (5–15 min each). Rehydration was performed through a graded ethanol series (100%, 95% and 75%) to distilled water, followed by thorough washing in water.

To enhance fibrin staining and colour differentiation, sections were mordanted in Bouin’s fixative at 56 °C for 1 h and subsequently rinsed extensively in running tap water to remove excess picric acid.

Immediately before use, Weigert’s iron haematoxylin working solution was prepared by mixing equal volumes of solutions A and B. Sections were stained with Weigert’s iron haematoxylin for 10 min, briefly rinsed in water, and differentiated in 1% acid alcohol until nuclei remained slightly overstained. Following washing in running tap water, sections were blued in Scott’s tap water substitute and rinsed again in running water.

Slides were then immersed in 95% ethanol before staining with Martius yellow solution for 5 min. After a brief wash in running tap water, sections were stained with crystal scarlet solution for 5 min and rinsed again in water. Differentiation was performed using phosphotungstic acid solution for 10 min, with additional incubation applied where required following microscopic assessment. Sections were subsequently washed in running tap water and counterstained with aniline blue solution for 5 min. Finally, sections were dehydrated through graded alcohols, cleared in xylene and mounted with a permanent mounting medium and coverslip. Slides were allowed to dry overnight before imaging by light microscopy.

Stained sections were imaged using a Leica THUNDER microscope equipped with 10×, 20×, 40×, 63× and 100× objective lenses. Representative regions of interest were acquired under identical acquisition settings within each experimental series to ensure consistency between samples. Images were captured and processed using the manufacturer’s standard acquisition software.

Whole-slide imaging was performed using a Hamamatsu slide scanner with DAPI and FITC fluorescence channels. Slides were scanned at 40× magnification using an exposure setting of 20 ms. Digital image files were quality checked following acquisition to confirm focus uniformity and channel alignment across the full tissue section.

18.	*Unknown article.*
